# Optimised in-solution enrichment of over a million ancient human SNPs

**DOI:** 10.1101/2024.05.16.594432

**Authors:** Roberta Davidson, Xavier Roca-Rada, Shyamsundar Ravishankar, Leonard Taufik, Christian Haarkötter, Evelyn Collen, Matthew Williams, Peter Webb, M. Irfan Mahmud, Erlin Novita Idje Djami, Gludhug A. Purnomo, Cristina Santos, Assumpció Malgosa, Linda R. Manzanilla, Ana Maria Silva, Sofia Tereso, Vítor Matos, Pedro C. Carvalho, Teresa Fernandes, Anne-France Maurer, João C. Teixeira, Raymond Tobler, Lars Fehren-Schmitz, Bastien Llamas

## Abstract

In-solution hybridisation enrichment of genetic markers is a method of choice in paleogenomic studies, where the DNA of interest is generally heavily fragmented and contaminated with environmental DNA, and where the retrieval of genetic data comparable between individuals is challenging. Here, we benchmarked the commercial “Twist Ancient DNA” reagent from Twist Biosciences using sequencing libraries from ancestrally diverse ancient human samples with low to high endogenous DNA content (0.1–44%). For each library, we tested one and two rounds of enrichment, and assessed performance compared to deep shotgun sequencing. We find that the “Twist Ancient DNA” assay provides robust enrichment of ∼1.2M target SNPs without introducing allelic bias that may interfere with downstream population genetics analyses. Additionally, we show that pooling up to 4 sequencing libraries and performing two rounds of enrichment is both reliable and cost-effective for libraries with less than 27% endogenous DNA content. Above 38% endogenous content, a maximum of one round of enrichment is recommended for cost-effectiveness and to preserve library complexity. In conclusion, we provide researchers in the field of human paleogenomics with a comprehensive understanding of the strengths and limitations of different sequencing and enrichment strategies, and our results offer practical guidance for optimising experimental protocols.

## Introduction

One of the major challenges faced when working with ancient DNA (aDNA) is the high proportion of exogenous DNA contamination present in the DNA extract. This contamination is primarily due to microbes invading the organism *post-mortem*, present in the soil where the specimen was buried, or introduced during sample handling and laboratory processes. To counteract this, a method that has become popular is the in-solution enrichment of target genomic regions using pre-designed oligonucleotides as molecular “probes” or “baits”.

Compared to shotgun sequencing, this technique increases the proportion of target DNA in a sequencing library, lowering sequencing costs required to produce adequate comparable data across individual samples.

In 2012, Patterson and colleagues proposed a molecular bait design for application in human paleogenomic research that made use of a particular ascertainment technique to enable population genetics studies of global human populations over time (Patterson et al., 2012). The bait design was ultimately restricted to approximately 1.2 million genome-wide SNPs, and became known as the ‘1240k reagent’, leading to the generation of thousands of individual genome-wide datasets (Marciniak & Perry, 2017; Olalde & Posth, 2020; Skoglund & Mathieson, 2018). However, since the original publication of the molecular baits sequences in 2015 (Fu et al., 2015; Haak et al., 2015; Mathieson et al., 2015), the legacy 1240k reagent has only been available through a commercial arrangement to a small number of research groups. This presented researchers with the choice of either collaborating with these groups to access the 1240k reagent or using the more expensive deep shotgun sequencing to obtain adequate data compatible with the 1240k SNP loci.

In 2021, two biotechnology firms, Daicel Arbor Biosciences and Twist Bioscience, produced commercial in-solution enrichment kits targeting the same 1240k SNPs plus, in each, an additional set of variants, and made these kits available to every research group (Rohland et al., 2022). However, recent studies have revealed a strong allelic technical bias in data generated with the Daicel Arbor Biosciences baits (Davidson et al., 2023; Rohland et al., 2022). While a comparatively mild allelic bias is also present in the legacy 1240k reagent (Davidson et al., 2023; Rohland et al., 2022), this has not previously been an issue as all enriched human paleogenomic data were generated with this legacy 1240k reagent and thus co-analysable.

While many researchers rely on enrichment methods to obtain affordable paleogenomic data, those wary of potential problems arising from the reported biases may prefer the more expensive shotgun sequencing approach. Indeed, because bait binding affinities differ for each allele at a targeted site, all target-hybridisation enrichment approaches are expected to have some allelic bias. Accordingly, an important consideration for researchers is whether the realised bias is strong enough to meaningfully affect population genomic analyses and interpretations. Furthermore, paleogenomic research depends upon the comparison of newly generated data to the cumulative set of published genome-wide datasets, making it essential that biases are not introduced when comparing data generated with different methods. Such biases may arise from differences in the assay design or when researchers use untested protocols that deviate from the manufacturer’s implementation. For example, some users may wish to pool libraries for input into a single enrichment reaction to decrease the reagent cost per library. However, since the enrichment protocol requires amplifying the libraries using PCR, pooling different libraries might increase instances of index-hopping between DNA molecules and introduce biases or cross-contamination between libraries co-enriched within the same reaction (Lahr & Katz, 2009; MacConaill et al., 2018; Meyerhans et al., 1990; Mitra et al., 2015; Sinha et al., 2017). Another example involves the choice to perform an extra round of enrichment to increase the target DNA yield, which will likely amplify any biases present in the first enrichment round (Davidson et al., 2023). While deviations from standard protocols are common, they can introduce variability into potential allelic biases that may be dangerous for downstream comparative analyses if not thoroughly tested and transparently reported.

Previous work by Rohland and colleagues found no evidence of allelic bias in the Twist assay (Rohland et al., 2022). However, their study was confined to well preserved West-Eurasian samples. Therefore, in this study, we aim to benchmark the commercial “Twist Ancient DNA” reagent from Twist Biosciences, using 24 ancient human samples, from four populations from three continents, across a range of endogenous DNA percentages. We compare deep shotgun sequencing, one and two rounds of enrichment with the Twist Bioscience “Twist Ancient DNA” reagent for cost-effectiveness and allelic biases. We also compare enrichment efficacy and biases between single and pooled library enrichments.

## Materials and Methods

### Facilities

Pre-amplification experiments were performed at the Australian Centre for Ancient DNA (ACAD)’s ultra-clean laboratory facilities following rigorous laboratory procedures to minimise contamination and ensure high standards of quality for the genetic data (Llamas, Valverde, et al., 2017; Llamas, Willerslev, et al., 2017). All post-amplification experiments were completed in standard molecular biology laboratories at the University of Adelaide and subsequent bioinformatics workflows executed on the University of Adelaide’s HPC.

### DNA extraction and Library Preparation

Skeletal remains from 24 ancient humans sourced from Iberia, Central America, and Southeast Asia were used for these experiments. Prior to DNA extraction, skeletal samples were sterilised using UV, bleach, and ethanol to minimise surface contamination. Approximately 0.1 g of bone powder was used for DNA extraction. Ancient DNA molecules were retrieved using a method optimised for degraded DNA (Dabney et al., 2013) and partially UDG-treated (Rohland et al., 2015) double-indexed double-stranded DNA libraries were subsequently generated (Meyer & Kircher, 2010). Quality control and quantification steps were completed using Qubit (Thermo Fisher) and TapeStation (Agilent) prior to over-amplification and enrichment or shotgun sequencing.

### Library Enrichment

Libraries were over-amplified in order to reach the 1000 ng needed for enrichment. For each library, the PCR reaction mix consisted of 5-10 µl of library, 25 µl of KAPA HiFi HotStart ReadyMix (Roche), 5µl each of 10 µM IS5 and IS6 primers (Rohland et al., 2015) and ultrapure water in a total volume of 50 µl. PCR amplification was performed with an initial denaturation and polymerase activation at 98°C for 2 min, 15 cycles of 98°C for 20 sec, 56°C for 30 sec, 72°C for 45 sec, and final extension at 72°C for 5 min. DNA purification was performed using 1.2x AmpureXP beads with two 80% ethanol washes, and the DNA was eluted in 30 µl of water.

To compare the performance and cost-effectiveness of single enriched libraries versus pooled enriched libraries, we prepared i) six reactions, each consisting of 1000 ng of DNA from a single library, each exhibiting varying endogenous DNA percentages as determined from prior screening shotgun sequencing runs; ii) three reactions consisting of two low endogenous DNA pooled libraries; and iii) three reactions consisting of four high endogenous DNA pooled libraries (Figure 1, Table S1). While each enrichment reaction contained a final quantity of 1000 ng of DNA, the amount of DNA required per library was reduced due to pooling, allowing for a decrease in PCR cycles to avoid overamplification and maintain library complexity. Library pooling was calculated from total DNA quantification of each library, rather than endogenous content calculated from the shotgun screening data. Moreover, pools of two libraries were configured to include Iberian and Central American or Southeast Asian samples, and pools of four were configured to include three samples of Iberian origin and the remaining sample having either Central American or Southeast Asian ancestry. This enabled the evaluation of cross-contamination created by dual index-hopping across DNA fragments with different ancestries in pooled library enrichments.

**Figure 1:**
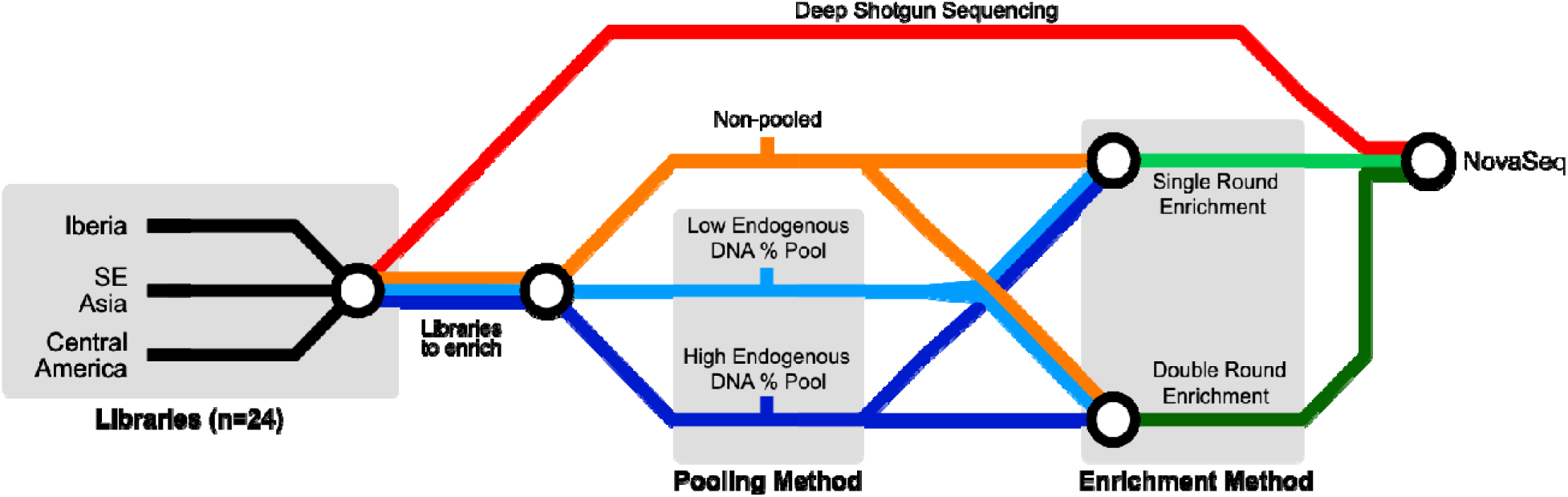
24 aDNA libraries, representing ancestries from Iberia, Southeast Asia, and Central America were selected, all underwent deep shotgun sequencing, and were also input into enrichment experiments. This included six unpooled libraries that were enriched individually, while the remaining libraries were pooled into six distinct reactions: 3 reactions with 2 low endogenous DNA libraries each, and 3 reactions with 4 high endogenous DNA libraries each. All sets of pooled and unpooled libraries underwent one as well as two rounds of enrichment prior to sequencing.

Enrichment was performed using the Twist Bioscience “Twist Ancient DNA” reagent following the manufacturer’s protocol. For each input pool, we independently performed one and two rounds of enrichment (henceforth labelled as TW1 and TW2, respectively) in order to compare performance and cost effectiveness. The post-enrichment PCR amplification was performed using KAPA HiFi HotStart ReadyMix (Roche) and IS5 and IS6 primers as described above, with a 98°C initialisation for 24 sec, 15 cycles (1st round) or 7 cycles (2nd round) of 98°C for 15 sec, 60°C for 30 sec, 72°C for 30 sec , and a 72°C final extension for 60 sec. DNA purification, and library quality control and quantification were performed as described above.

After one round of enrichment, each library underwent a reconditioning PCR to reduce heteroduplexes: libraries were concentrated down to 5 µl and mixed with 10 µl of Herculase Buffer (Agilent), 5 µl of 2.5 nM dNTPs, 1 U of Herculase II Fusion (Agilent), 1 µl each of 10 µM IS5 and IS6 primers and ultrapure water in a final volume of 50 µl, then reconditioned with one cycle of 95°C for 2 min, 58°C for 2 min, and 72°C for 5 min. Where possible, a lower number of post-enrichment amplification cycles were performed (e.g., 7–10 cycles) in order to avoid heteroduplexes, which depended on the initial endogenous DNA percentage. DNA purification, and library quality control and quantification were performed as described above.

### Sequencing

All shotgun and enriched libraries were sent for sequencing using a NovaSeq 6000 System with a 2 × 100 bp SP Flow Cell in XP mode at the Kinghorn Centre for Clinical Genomics (Sydney, NSW, Australia).

### Data processing

Raw data were processed with the aDNA analysis workflow package nf-core/eager version 2.4.6 (Fellows Yates et al. 2021). Merged read mates were mapped to the GRCh37d5 reference genome using bwa aln with parameters -l 1024 -n 0.01 -o 2 (Oliva et al., 2021). Two nt were trimmed from the terminal ends of all retained reads using the trimBam function of bamUtil (https://github.com/statgen/bamUtil). Standard quality filters (mapping quality ≥ q25 and base quality ≥ Q30) were applied through samtools version v1.12 mpileup function (Danecek et al., 2021). Reads were deduplicated using MarkDuplicates from Picard. Pseudohaploid variant calling using the Twist Bioscience “Twist Ancient DNA” SNP panel (Rohland et al., 2022) was performed using pileupCaller (https://github.com/stschiff/sequenceTools).

Ancient DNA authenticity, endogenous DNA percentage, fragment size distribution, and post-mortem damage rate at the read termini were determined using DamageProfiler (Neukamm et al., 2021).

### Mitochondrial data processing

Raw data were processed with the aDNA analysis workflow package nf-core/eager version 2.4.6 (Fellows Yates et al., 2021). Merged reads with a length greater than or equal to 30 nt were mapped to the mitochondrial revised Cambridge Reference Sequence (rCRS) using CircularMapper (https://github.com/apeltzer/CircularMapper) and bwa aln with parameters -l 1024 -n 0.01 -o 2 -k 2 (Oliva et al., 2021). Read trimming and filtering followed the procedures outlined above. The read pileups were visually inspected in Geneious v2022.1.1 (Biomatters; https://www.geneious.com). Mitochondrial contamination estimates were calculated using mitoverse HaploCheck version 1.3.2 (Weissensteiner et al., 2021).

### Assessment of enrichment efficacy

We note that the word “endogenous”, in the term “endogenous DNA content” is loosely interpreted in the paleogenomic literature and often refers to different quantifications. Here, we consider endogenous DNA as the DNA mapping to the reference genome of interest, whether that DNA pertains to the sample or to contaminant sources. Hereafter, we refer to endogenous DNA percentage as “endo%”. We propose the following nomenclature for different calculations of endo%, regarding the specific read counts in the calculation, and whether the calculation is applied to shotgun sequencing data or enriched data. Importantly, we distinguish ” endo%”, referring to the state of DNA found in the sample, from “post-enrichment endo%” referring to the enriched DNA from the taxon of interest after in-solution target enrichment. For both of these, we define calculations for the sequenced, mappable, filtered and unique endo% as each informs differently about the usability of the sample and sequenced data. See Tables 1 and 2 for further information.

**Table 1:** Definitions of different read counts.

| Read count name | Definition |
| --- | --- |
| Sequenced | Number of sequenced reads (i.e paired-end read pairs) in the demultiplexed fastq file before filtering or collapsing |
| Pre-mapping | Number of reads input to mapping after collapsing, read length and base quality filtering |
| Mapped | Number of reads mapping to the target reference genome before further filtering |
| Filtered | Number of reads mapping to the target reference genome with mapping quality $\geq 25$ before further filtering |
| Dedup | Number of reads retained after filtering mapped reads and removing PCR duplicates |

**Table 2:** Definitions of different quantifications of endogenous DNA percentage in sequencing libraries.

| Definition | Data | Formula (x100) | Read length filter | Mapping quality filter | Deduplication | Purpose | Comments |
| --- | --- | --- | --- | --- | --- | --- | --- |
| Sequenced endo% | Shotgun | Mapped | Y | N | N | Informs about sample preservation and need for sample enrichment | Estimates are the closest to the true biological endogenous DNA content |
|  |  | Sequenced | N | N | N |  |  |
| Mappable endo% | Shotgun | Mapped | Y | N | N | Informs about sample preservation and need for sample enrichment | Most commonly used in paleogenomic research, but may inflate endogenous DNA estimates when very short reads are abundant |
|  |  | Pre-mapped | Y | N | N |  |  |
| Filtered endo% | Shotgun | Filtered | Y | Y | N | Informs about sample preservation and need for sample enrichment and data robustness | More informative than the above for reads used in downstream analyses |
|  |  | Pre-mapped | Y | N | N |  |  |
| Dedup endo% | Shotgun | Dedup | Y | Y | Y | Informs about sample enrichment, sample preservation, data robustness, and library complexity | Indicator of library complexity, provides the “useful” proportion of shotgun sequencing data for analyses. Consider in decision to enrich as high duplication, when amplified, can overwhelm unique molecules |
|  |  | Pre-mapped | Y | N | N |  |  |
| Sequenced post-enrichment endo% | Enriched | Mapped | Y | N | N | Informs about enrichment efficacy | n.a. |
|  |  | Sequenced | N | N | N |  |  |
| Mappable post-enrichment endo% | Enriched | Mapped | Y | N | N | Informs about enrichment efficacy | n.a. |
|  |  | Pre-mapped | Y | N | N |  |  |
| Filtered post-enrichment endo% | Enriched | Filtered | Y | Y | N | Informs about enrichment efficacy and data robustness | n.a. |
|  |  | Pre-mapped | Y | N | N |  |  |
| Dedup post-enrichment endo% | Enriched | Dedup | Y | Y | Y | Informs about enrichment efficacy, data robustness, and | Provides the “useful” proportion of enriched data for |
|  |  | Pre-mapped | Y | N | N |  |  |
|  |  |  |  |  |  | library complexity retention | analyses |

### Cost-benefit analyses

We calculated the financial cost of the experiment on a per SNP basis using the equation below, which captures the combined costs incurred by enrichment and DNA sequencing. The relative difference between the cost of enrichment and the cost of sequencing is expected to impact the economic benefits of enrichment on a case-by-case basis, depending on laboratory-specific commercial agreements for reagents purchase and sequencing service provision, and the sequencing effort (red variables in the equation).

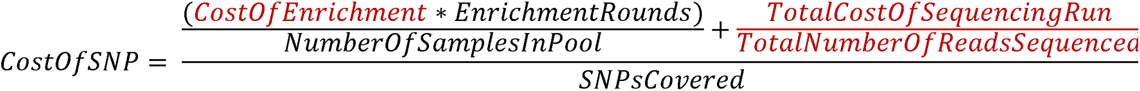

### Allelic bias

Allelic bias was measured using *f*_4_ statistics calculated using *qpDstat* v980 in *f*_4_ mode with options *f4mode: YES* and *inbreed: YES* in AdmixTools v. 7.0.2 (Patterson et al., 2012).

### Index hopping

To estimate single-index hopping, we demultiplexed all possible index combinations of the double-indexed libraries. Reads that resulted from the combination of indices initially assigned to two different samples were considered to be the result of index hopping. Subsequently, the rate of single-index hopping was determined for every pairwise combination of samples, expressed as the ratio of hopped reads to the sum of hopped reads and all retrievable reads from a sample pair.

For libraries enriched in pools, separate contamination estimates were obtained for reads mapping to the Y chromosome, autosomes and mitochondrial genome and were used to estimate the impact of double-index hopping. We used Haplocheck v1.3.2 (Weissensteiner et al., 2021) to estimate mitochondrial contamination for all the sequenced libraries. The genetic sex of the samples was determined based on the results of SexDetERRmine (https://github.com/nf-core/modules/tree/master/modules/nf-core/sexdeterrmine) from the deep shotgun sequencing data. For all the male samples we used HapConX (Huang & Ringbauer, 2022), an aDNA contamination estimation tool that works using a haplotype copying framework for male X chromosomes to estimate cross-contamination between samples, which in this case is equivalent to double-index hopping. Similarly, we calculated contamination estimates using ANGSD (Korneliussen et al., 2014; Rasmussen et al., 2011) for all samples, including the females as a positive control.

Finally, we ran DICE (Racimo et al., 2016), a Bayesian method to estimate the rate of contamination from a specific ancestry, in two-population mode to calculate nuclear contamination for the Central American sample in the pool of four, the three other samples bearing Iberian ancestry. The contaminating population was set to Iberia (IBS) and the anchor population was set to an appropriate Central American population from the 1000 Genomes dataset. The program was run with a Markov chain of 1 million steps and the contamination estimate was restricted to 100 nt either side of each targeted 1240k SNP. We tried to run DICE for Southeast Asian samples with the KHV population from the 1000 Genomes dataset as the anchor. However, the combination of high genetic diversity and underrepresentation of Southeast Asians in the 1000 Genomes dataset meant that we were unable to obtain cross contamination estimates for our Southeast Asian samples using DICE.

## Results

### Method comparison

In human paleogenomics, it is common practice to shotgun sequence libraries at a low depth to obtain “screening” data and gather information about library quality (e.g., complexity, endo%) in order to make decisions about further processing. Depending on sample quality and budget, this may lead to deeper shotgun sequencing, target enrichment, or discarding the sample from the experiment. In this study, we selected 24 samples based on the mappable endo% calculated from shotgun screening sequencing data, with these samples also submitted to deeper shotgun sequencing for the purpose of this study. We subsequently compared read data obtained from the shallow screening and deep shotgun sequencing steps to better understand how well shotgun screening data reflects the true quality of the DNA sequencing library (Figure S1). For all measures of SNP count and endo%, the shotgun screening and deep shotgun results have a strong positive linear correlation with high *r*^2^ values (0.69–0.95). According to the fitted trendlines, deep shotgun sequencing retrieves slightly less SNPs per million reads (slope = 0.86), and a slightly higher endo% than suggested by the screening data (slope = 1.02) (Figure S1). These indications are based on the data from only 24 libraries. However, we expect that a larger sample size would likely show the same trends, keeping in mind that samples will likely exhibit inconsistent individual variation between screening and deeper shotgun sequencing.

Comparing the efficacy of deep shotgun sequencing to one and two rounds of enrichment (TW1 and TW2, respectively), we observe that TW2 consistently captured more SNPs per sample in our experiments (Figure 2A). However, when normalising the data per million sequenced paired reads (Figure 2B), reads into mapping (Figure 2C), mapped reads (Figure 2D), mapping quality filtered reads (Figure 2E), or deduplicated reads (Figure 2F), at least 3 out of 4 libraries with mappable endo% > 38% produced less SNPs per million reads after a second round of enrichment. Although two rounds of enrichment consistently yields higher sequenced, mappable, and filtered post-enrichment endo% (Figure 2G–I), the unique post-enrichment endo% was higher only for libraries with mappable endo% < 38% (Figure 2J). Overall, our results show that two rounds of enrichment may be detrimental to SNP yield for high quality libraries. Furthermore, our results highlight the impact of the method used to calculate endo% in the context of enrichments, especially for high endo% libraries.

**Figure 2:**
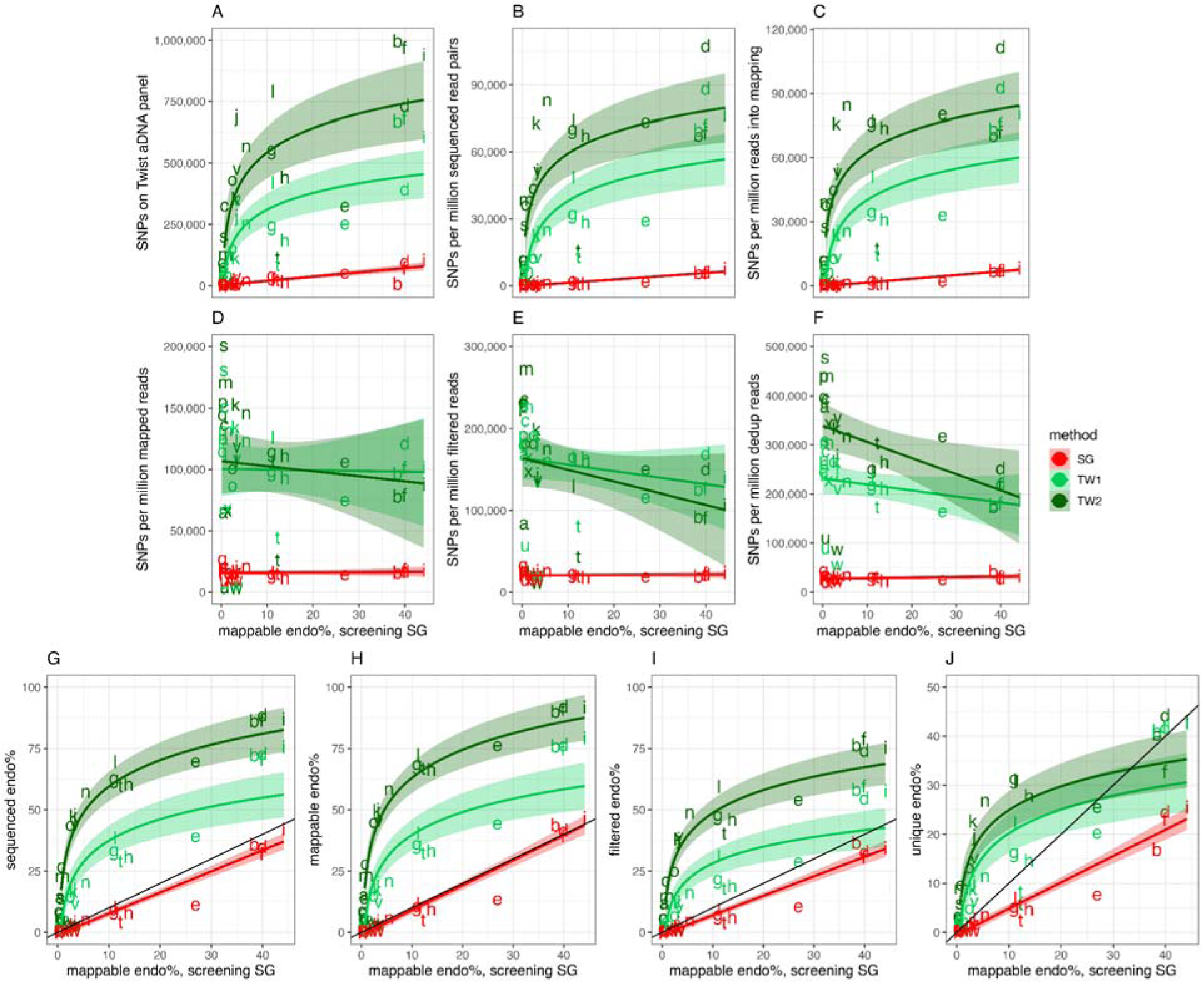
Enrichment efficacy of deep shotgun sequencing or Twist enrichment (either one or two rounds, i.e., TW1 or TW2, respectively) in relation to the mappable endo% from the screening shotgun data (x-axis). Enrichment efficacy was measured using (A) total on-target SNPs per sample, (B) SNPs per million sequenced read pairs, (C) SNPs per million reads going into mapping, (D) SNPs per million mapped reads, (E) SNPs per million filtered reads, passing filter for MAQ>25, (F) SNPs per million unique reads, (G) sequenced post-enrichment endo%, (H) mappable post-enrichment endo%, (I) filtered post-enrichment endo%, and (J) unique post-enrichment endo%. Plotted letter points are consistent between the same sample. The solid lines and shaded areas show fitted linear or logarithmic transformed linear regression models as appropriate. The black line in panels G–J shows y = x.

### Cost effectiveness

One key value of performing library enrichment is the potentially significant reduction in sequencing cost to generate useful data. We tested the cost effectiveness of deep shotgun sequencing and Twist enrichment using one or two enrichment rounds by applying all three approaches on the same set of 24 samples. While the specific costs are unique to our experiment, it is clear that the Twist enrichment (1 or 2 rounds) is more cost-effective per SNP than shotgun sequencing. This advantage is greatest for libraries with low mappable endo%. Further, our fitted logarithmic model predicts that 2 rounds of enrichment was more cost-effective per SNP than 1 round only for libraries with mappable endo% < 20% (Figure 3). We note, however, that this threshold depends on the relative costs of enrichment and sequencing and thus should be considered as a guide rather than a definitive value for future studies. We also compared the relationship between cost per SNP and the number of retrieved target SNPs (Figure 4), which revealed that Twist enrichment obtained more total and relative target SNPs than shotgun sequencing. Additionally, for the majority of samples, 2 rounds of enrichment generated more SNPs and was more cost-effective than a single round (Figure 4). We note that the cost of enrichment and sequencing might vary significantly depending on pricing agreements with manufacturers, distributors, and service providers, and we encourage other research groups to carefully consider their own circumstances when determining the number of rounds of enrichment for their study.

**Figure 3:**
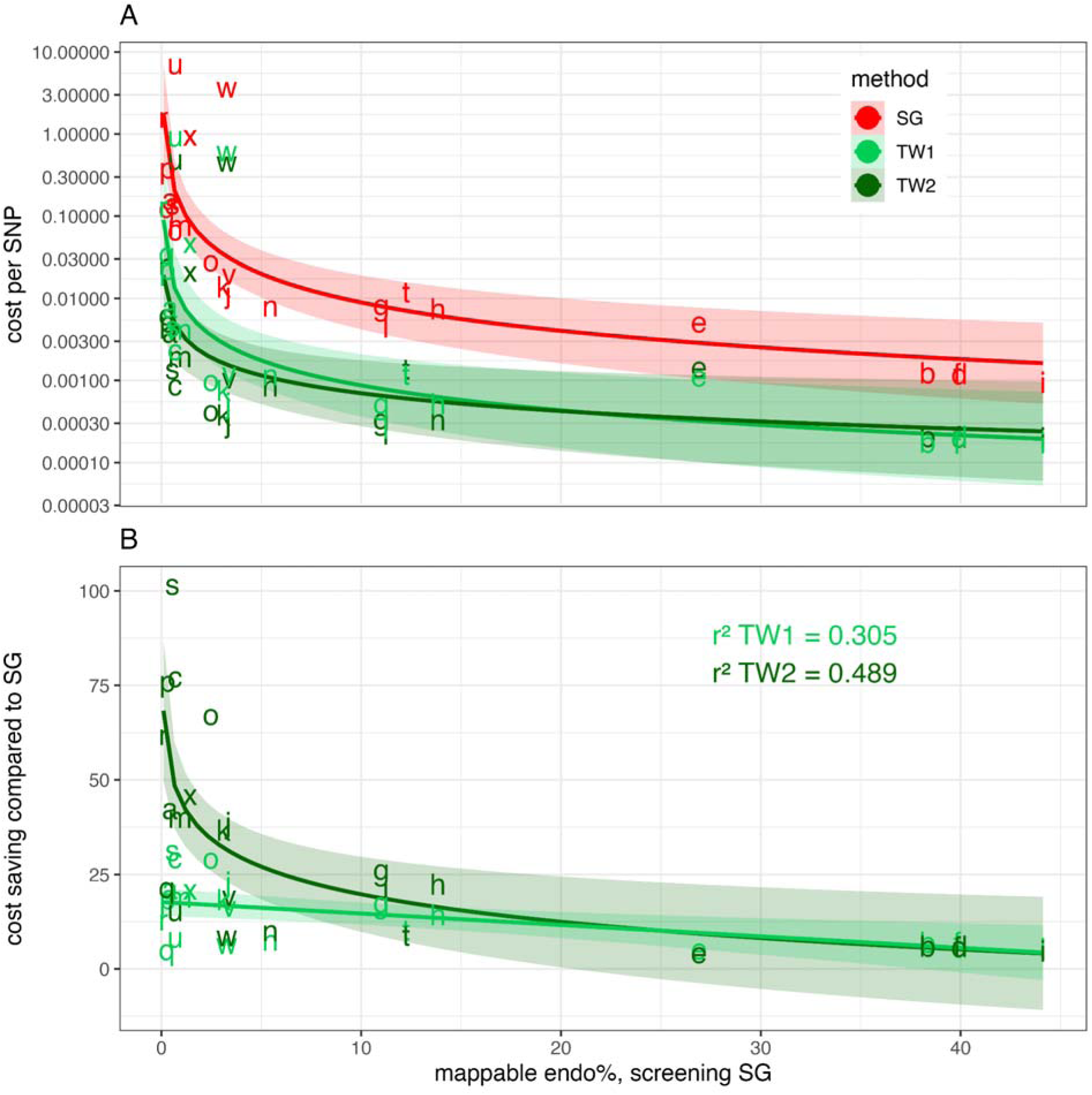
(A) Cost per SNP (AUD) obtained from shotgun sequencing and Twist enrichment methods as a function of the mappable endo%. The y-axis is log10 transformed. (B) Relative fold cost saving per SNP (AUD) obtained from TW1 and TW2 compared to deep shotgun sequencing. Plotted letter points are consistent between the same sample. Linear or log10 transformed linear models are fit to each method (solid lines), with 95% confidence intervals (shaded areas). This figure is also available as an interactive app where users can input their own laboratory costs. See supplementary information.

**Figure 4:**
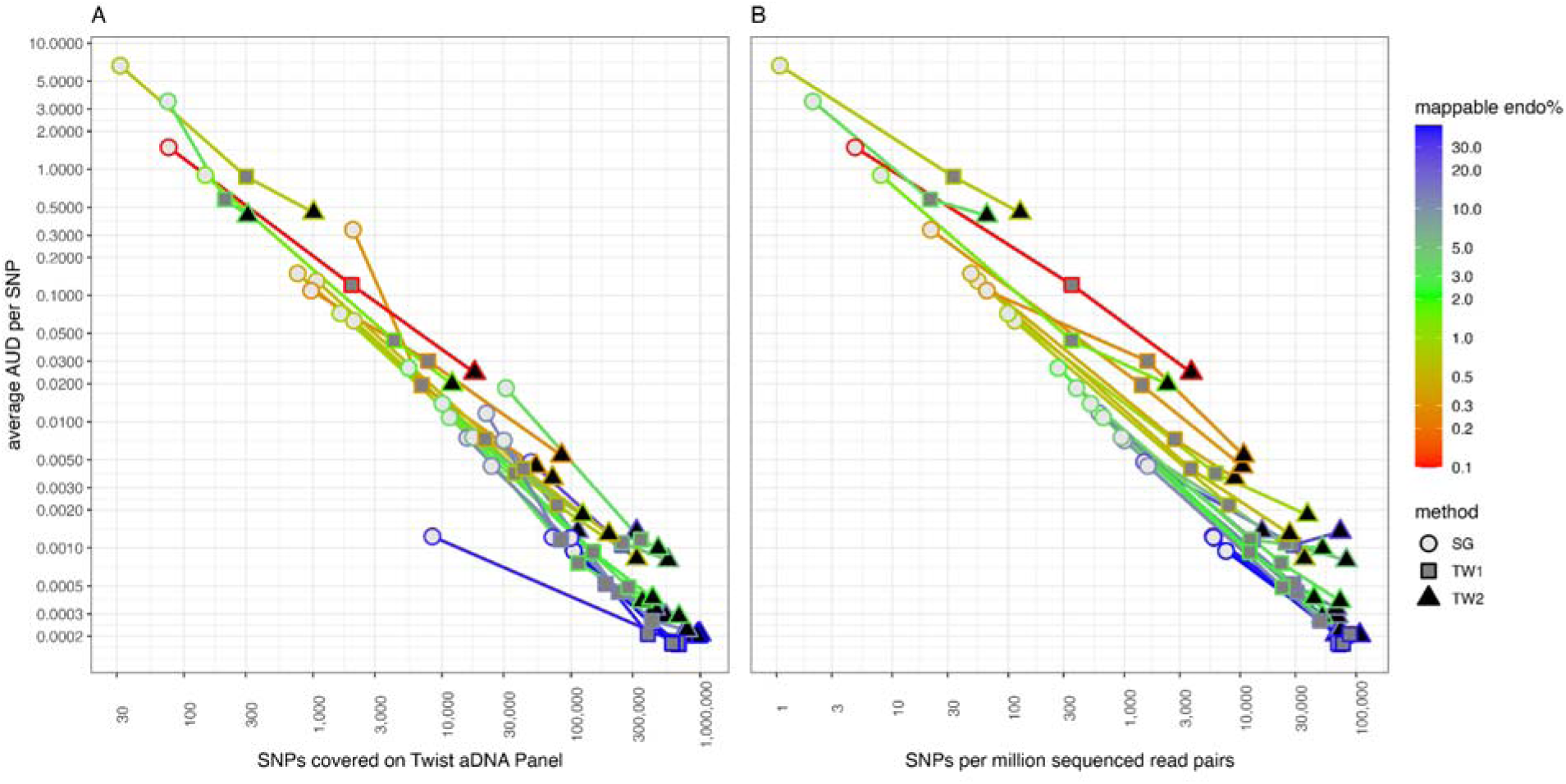
Cost per SNP in relation to SNP coverage (x-axis) and mappable endo % measured from screening shotgun data (colours) for deep shotgun sequencing and enrichment methods (shapes). (A) Total SNP coverage on Twist ancient DNA panel and (B) SNPs per million sequenced read pairs. Lines connect the same library across the three methods. Plot axes and colour scale are log10 transformed.

### Allelic bias

To measure allelic bias we performed *f*_4_ statistics of the form *f*_4_ (Mbuti, Pop1.MethodA; Pop2.MethodA, Pop2.MethodB), where the populations were one of East Iberia, West Iberia, Central America or Southeast Asia and the methods were one of shotgun, TW1, or TW2. Given that the same population is present in the right hand pairing of the equation and an outgroup to Eurasian and American populations placed in the first position, significant deviations from *f*_4_ = 0 are indicative of allelic bias between the tested Pop1.MethodA left population and one of the Pop2 right populations. The bias may signal differential allelic affinities across the assay methods (Figure 5) (Davidson et al., 2023).

**Figure 5:**
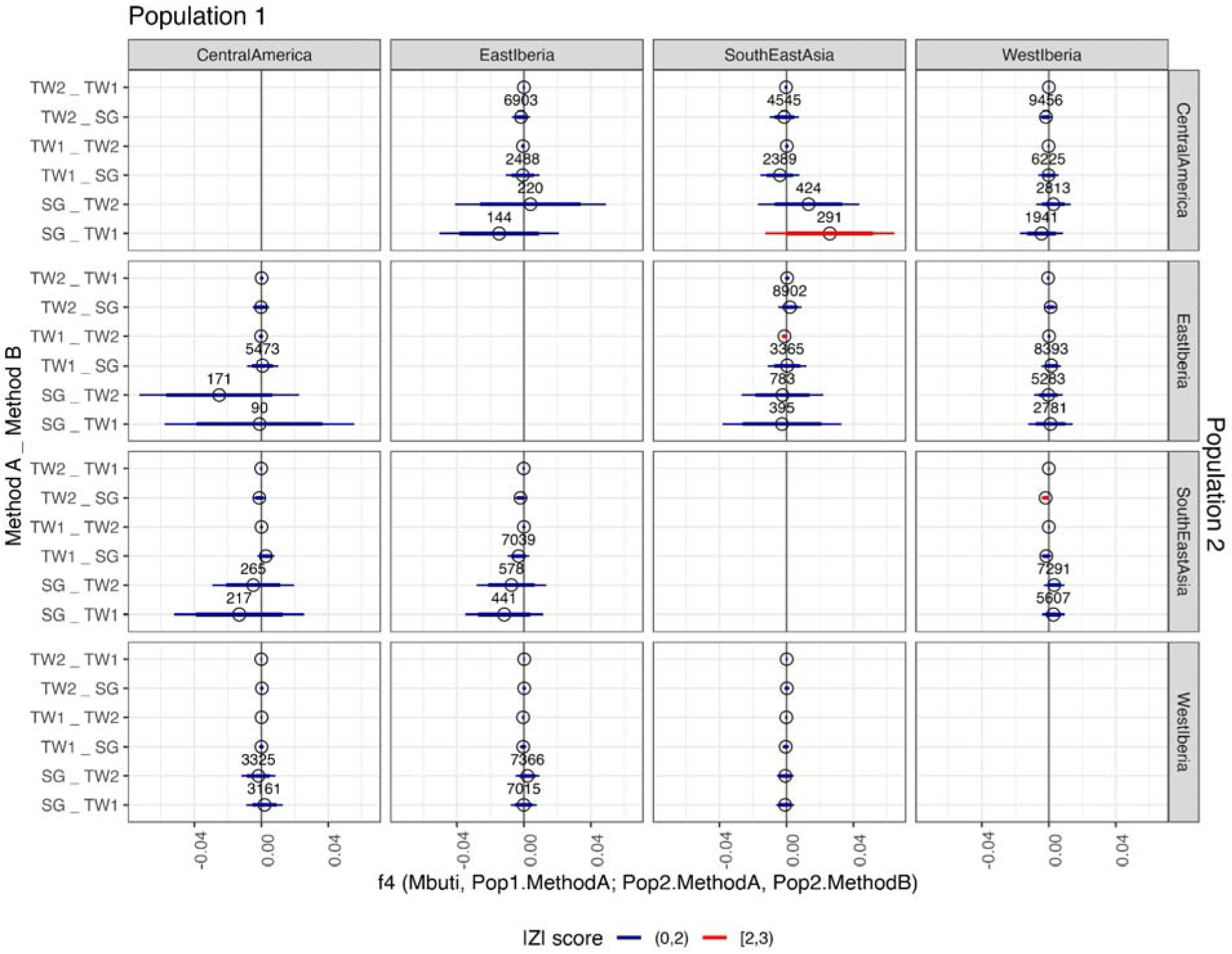
*f*_4_ statistics of the form *f*_4_(Mbuti, Pop1.MethodA; Pop2.MethodA, Pop2.MethodB). Populations 1 and 2 are faceted horizontally and vertically, respectively. Thick and thin error bars represent 2 and 3 s.e. respectively. |Z| > 2 are coloured red, no test has |Z| > 3. Tests with < 10,000 SNPs are annotated with the number of SNPs.

In all tests, the absolute Z score is less than 3, and the majority are less than 2, indicating that none of the observed *f*_4_ statistics are significantly different from zero. Accordingly, we find no evidence for significant allelic assay bias from the Twist enrichment, whether between one and two rounds of enrichment, or between any enrichment and shotgun data, providing support for the combined analysis of Twist and whole genome sequencing data (Figure 5).

### Impact of pooling

A concern when pooling libraries for enrichment is that adding more libraries into the reaction may decrease the yield from each individual library. To evaluate this, we compared the number of SNPs obtained per million reads against the mappable endo% to see if pools with fewer libraries generated more SNPs per library (Figure S3). Point distributions for the three types of pool overlap and linear trendlines for pooled libraries suggest higher SNP yield than in unpooled libraries.

Another unknown regarding the pooling of libraries for enrichment is how equitably the enrichment reaction captures targeted SNPs from each of the libraries, and if any inequities would be amplified by two rounds of enrichment. To investigate this, we compared various read counts from successive stages of data processing, between each library in every pool (Figure 6). The proportion of the total reads contributed by each library in a pool remains quite consistent between TW1 and TW2. In most pools, the read proportions are similar across the different read counts, with the biggest observable change being between the reads into mapping and mapped reads when mappable endo% is low (Figure 6). This is likely because the pooling calculations are based on DNA concentration of the library rather than the endo% of each library. Of note, the high mappable endo% library in the HE4 pool that drops out after mapping (Figure 6, bottom sub-bar in pink in HE4 facets) is characterised by a high rate of duplication.

**Figure 6:**
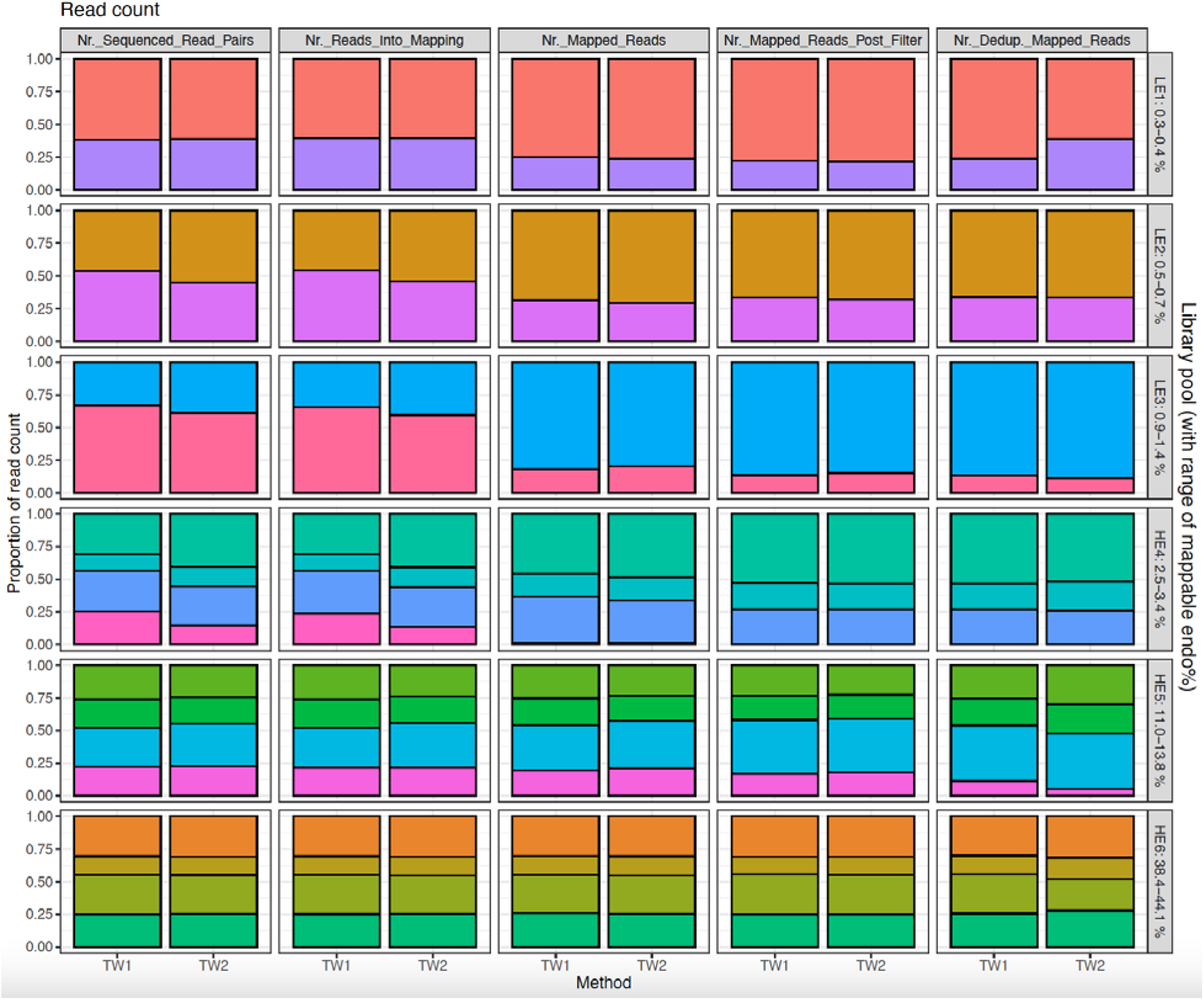
Equity of library enrichment in pooled reactions. Library pools are ordered from low to high endo% from top to bottom and labelled with the range of mappable endo%. Colours represent individual libraries in each pool. Panels show from left to right: number of sequenced read pairs, number of reads input to mapping, number of reads mapped, number of mapped reads passing mapping quality (≥ q25), and de-duplicated mapped reads.

Another concern when pooling libraries is “cross-contamination” of samples due to dual index hopping between DNA molecules from different libraries. Therefore, we first calculated the pairwise rate of single index hopping by counting all possible combinations of the library indexes included in our experiment. These pairwise hopping rates were evaluated separately for unpooled libraries and each pool size (2 and 4 libraries), knowing that index hopping observed in unpooled libraries can only occur during sequencing. We observe a significant difference between the single-index hopping rate for pooled and unpooled libraries (Figure 7), with no significant difference between one and two rounds of enrichment (Figure S2). Taking the observed mean pairwise single index hopping rate of ∼0.013 for pairs in 2-library pools (Figure 7), we estimate the expected proportion of dual index hopped reads is ∼0.00016, or 160 reads per million, assuming that indexes hop independently. Then, following the same assumptions, the observed mean pairwise single index hopping rate between a pair of libraries in a 4-library pool is ∼0.0059 (Figure 7), and given that there are six pairs of libraries in a 4-library pool, the estimated proportion of dual index hopped reads between any two libraries in a 4-library pool is ∼0.0059^2^ x 6 = ∼0.00021, or 208 reads per million.

**Figure 7:**
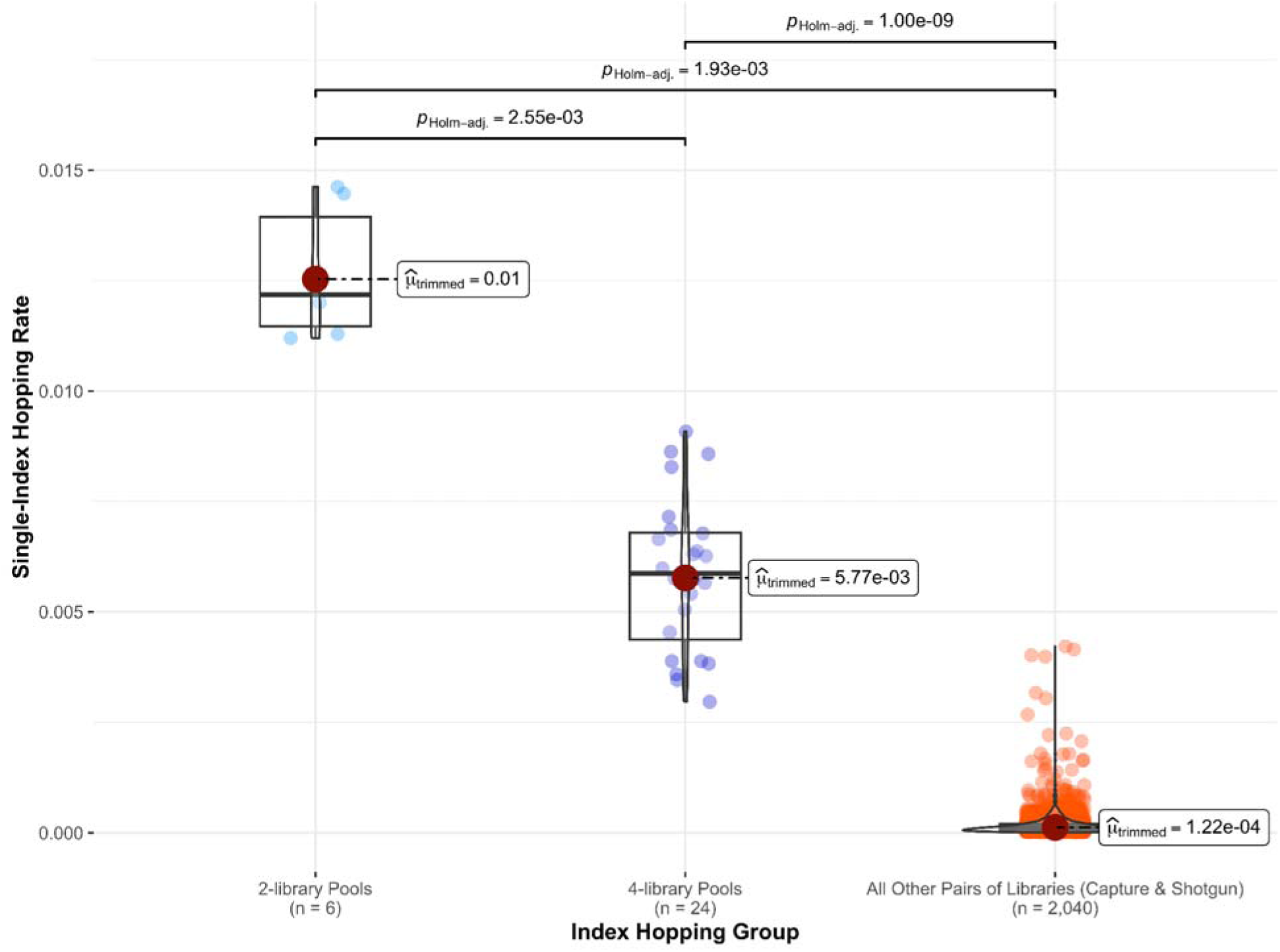
Pairwise rate of single index hopping between every pair of libraries, categorised into (left to right) 2-library pools, 4-library pools and all other pairs of libraries. Each category is annotated with the number of pairs (*n*). The hopping rate was calculated as the proportion of reads with a hopped index combination over the sum of reads from every possible index combination within the pair.

Furthermore, to investigate the prevalence of dual index hopping in our experiment we applied several tests for sample cross contamination, given that a dual index hopping event would appear in the sequencing data as contamination from another sample. We used HaploCheck to measure mitochondrial contamination, ANGSD and HapConX to measure contamination of the X chromosome in males and DICE to estimate autosomal contamination, with all results reported in Table S3.

Results from HaploCheck revealed the absence of detectable contamination in any sample (Table S3). However, five samples were reported with artifactual “heteroplasmies” by HaploCheck, raising the possibility of index-hopping contamination. Among the affected libraries, three (TW024_NA_TW2, TW015_HE6_TW2, and TW016_HE6_TW2) had fewer than 1000 reads mapping to the mitochondrial genome, presenting a challenge for haplogroup assignment. Notably, despite all three libraries being subjected to both single and dual rounds of enrichment, heteroplasmy was not detected in TW1. Of the remaining two affected samples, one library (TW006_HE6) was impacted in both TW1 and TW2, while the other (TW008_LE1) was once again only impacted in TW2. None of these artificial heteroplasmy loci were considered in haplogroup determination and when read pileups were inspected visually, they did not present a challenge for consensus sequence calling. Interestingly, we noticed unexpectedly low coverage for mitogenomes in Twist-enriched data, suggesting that increasing the number of mitochondrial probes during enrichment would be beneficial. Additionally, there was a decrease in mitochondrial genome coverage in TW2, indicating fewer unique reads and decreased complexity. Consequently, minimising PCR cycles is recommended in both the pre-enrichment over-amplification and enrichment PCRs.

X-chromosome contamination was estimated in the 18 male libraries in our experiment with ANGSD and HapConX. In the ANGSD results, all pools exhibit zero contamination for male samples whereas female samples are reported as contaminated, which is expected given the presence of two different X chromosomes and serves as a positive control. We caution that some contamination values reported by ANGSD are for samples with a total absence of X chromosome SNPs and are therefore ignored. HapConX reported extremely low estimated contamination for all samples, the highest being ∼0.037. Lastly, we ran DICE to estimate autosomal contamination based on ancestries, however this was only achievable for one pool (HE6; having one Central American individual and three Iberian individuals) due to poor data quality of low-endogenous pools, and a lack of relevant publicly available reference populations for our Southeast Asian samples. The DICE output converged upon a low contamination estimate of ∼4.55e-05 and 4.99e-04 for TW1 and TW2, respectively (Figure S3).

## Discussion

The findings of this study offer valuable insights into optimising enrichment strategies in human paleogenomics through the use of shotgun screening and pooling strategies for targeted enrichment. The positive linear relationship observed between shallow and deep shotgun sequencing data for SNP count and endo% underscores the reliability of shallow shotgun screening as a preliminary assessment tool for downstream analyses and decisions concerning aDNA library enrichment.

The high cost of deep shotgun sequencing compared to enrichment methods in contemporary paleogenomics is an important factor in favouring the latter approach, whether through the application of one or two rounds of enrichment. Our analyses show that, in general, it is most cost-effective to perform two rounds of enrichment, though a decrease in SNPs per million sequenced reads observed in high endo% libraries subjected to a second round of enrichment suggests that the benefits of further enrichment are dependent on sample quality. Comparing mappable and unique enriched endo% calculations shows that two rounds of enrichment for high mappable endo% libraries may result in sequencing a majority of duplicate reads, reducing the proportion of unique reads sequenced as well as the corresponding target SNP yield and mitochondrial genome coverage. This highlights the need for reducing the number of PCR cycles and/or opting for a single round of enrichment for libraries with high endo%—our study suggests the latter applies to libraries with mappable endo% > 38%, possibly > 27% (our experimental design did not include libraries with mappable endo% in the range 27%–38%).

Our assessment of allelic bias using *f*_4_ statistics yielded reassuring results, suggesting the absence of observable assay bias introduced by the Twist Bioscience “Twist Ancient DNA” reagent. This finding further consolidates the reliability of this reagent in producing unbiased results for paleogenomic analyses, providing a notable improvement compared to previously reported paleogenomic enrichment data generated with the legacy 1240k reagent and the Daicel Arbor Biosciences Expert Human Affinities Prime Plus enrichment kit (Davidson et al., 2023; Rohland et al., 2022).

Our exploration of the effects of pooling several libraries into a single enrichment reaction also yielded reassuring outcomes. We suggest that pooling up to four libraries does not have a substantial impact on SNP yield compared to single library reactions. We investigated the potential for cross-contamination due to dual index hopping between molecules originating from different libraries. Even though there is a significant difference in single-index hopping rates between pooled and unpooled libraries, the undetectability of dual-index hopping through the calculation of contamination estimates, coupled with its extremely low estimated occurrence rates, underscores the reliability and cost-effectiveness of the pooling approach.

## Acknowledgements

The work was supported by an Australian Research Council Discovery Early Career Researcher Award (DE190101069), the Australian Research Council Centre of Excellence for Australian Biodiversity and Heritage (CE170100015) and Bioplatform Australia. Laboratory work was conducted at The University of Adelaide with support from technical officer Navdeep Kaur. Computational analyses were conducted using supercomputing resources provided by the Phoenix HPC service at the University of Adelaide. We thank the Research Center for Prehistoric and Historic Archaeology, National Research Institute Agency and The Research Organization for Archaeology, Language, and Letters, National Research Institute Agency for providing the samples used for the analysis in this paper. All Southeast Asian samples used in the analysis are owned by their respective provider and by the communities in the region they were found. C.H. acknowledges the Spanish Ministry of Universities for funding the development of his PhD (FPU 20/01967). A.M.S., S.T., T.F., and V.M. acknowledge CIAS - Research Centre For Anthropology and Health, University of Coimbra, Portugal, funded by National Funds through the FCT - Foundation for Science and Technology, I.P., within the scope of the project UIDB/00283/2020 (https://doi.org/10.54499/UIDB/00283/2020). X.R.R. acknowledges the FCT - Foundation for Science and Technology, I.P./MCTES (PTDC/HAR-ARQ/6273/2020) for funding the development of his postdoctoral fellowship through the Portuguese National Funds (PIDDAC).

## Data Accessibility Statement

Individual de-identified genotype data will be made available upon conditional approval by the corresponding authors. Metadata including sequencing statistics and all analysis and plotting code is deposited here https://github.com/roberta-davidson/Davidson_etal_2024-Twist/.

## Author Contributions

Conceptualisation - RD, XRR, BL, MW, LFS

Lab - XRR, LT, CH, LFS, PW, JCT, MW, EC

Data Processing - SR, LT

Data Analysis - SR, RD, LT, XRR

Manuscript writing - LT, XRR, RD, BL, SR, RT

Manuscript editing – All authors

Funding - RT, BL

Sample Provision - MIM, ENID, CS, AM, LRM, AMS, ST, VM, JCT, PCC, AFM, TF

## References

Dabney, J., Knapp, M., Glocke, I., Gansauge, M.-T., Weihmann, A., Nickel, B., Valdiosera, C., García, N., Pääbo, S., Arsuaga, J.-L., & Meyer, M. (2013). Complete mitochondrial genome sequence of a Middle Pleistocene cave bear reconstructed from ultrashort DNA fragments. Proceedings of the National Academy of Sciences of the United States of America, 110(39), 15758–15763.

Danecek, P., Bonfield, J. K., Liddle, J., Marshall, J., Ohan, V., Pollard, M. O., Whitwham, A., Keane, T., McCarthy, S. A., Davies, R. M., & Li, H. (2021). Twelve years of SAMtools and BCFtools. GigaScience, 10(2). 10.1093/gigascience/giab008

Davidson, R., Williams, M. P., Roca-Rada, X., Kassadjikova, K., Tobler, R., Fehren-Schmitz, L., & Llamas, B. (2023). Allelic bias when performing in-solution enrichment of ancient human DNA. Molecular Ecology Resources, 23(8), 1823–1840.

Fellows Yates, J. A., Lamnidis, T. C., Borry, M., Andrades Valtueña, A., Fagernäs, Z., Clayton, S., Garcia, M. U., Neukamm, J., & Peltzer, A. (2021). Reproducible, portable, and efficient ancient genome reconstruction with nf-core/eager. PeerJ, 9(e10947), e10947.

Fu, Q., Hajdinjak, M., Moldovan, O. T., Constantin, S., Mallick, S., Skoglund, P., Patterson, N., Rohland, N., Lazaridis, I., Nickel, B., Viola, B., Prüfer, K., Meyer, M., Kelso, J., Reich, D., & Pääbo, S. (2015). An early modern human from Romania with a recent Neanderthal ancestor. Nature, 524(7564), 216–219.

Haak, W., Lazaridis, I., Patterson, N., Rohland, N., Mallick, S., Llamas, B., Brandt, G., Nordenfelt, S., Harney, E., & Stewardson, K. (2015). Massive migration from the steppe was a source for Indo-European languages in Europe. Nature, 522(7555), 207.

Huang, Y., & Ringbauer, H. (2022). hapCon: estimating contamination of ancient genomes by copying from reference haplotypes. Bioinformatics , 38(15), 3768–3777.

Korneliussen, T. S., Albrechtsen, A., & Nielsen, R. (2014). ANGSD: Analysis of Next Generation Sequencing Data. BMC Bioinformatics, 15(1), 356.

Lahr, D. J. G., & Katz, L. A. (2009). Reducing the impact of PCR-mediated recombination in molecular evolution and environmental studies using a new-generation high-fidelity DNA polymerase. BioTechniques, 47(4), 857–866.

Llamas, B., Valverde, G., Fehren-Schmitz, L., Weyrich, L. S., Cooper, A., & Haak, W. (2017). From the field to the laboratory: Controlling DNA contamination in human ancient DNA research in the high-throughput sequencing era. STAR: Science & Technology of Archaeological Research, 3(1), 1–14.

Llamas, B., Willerslev, E., & Orlando, L. (2017). Human evolution: a tale from ancient genomes. Philosophical Transactions of the Royal Society of London. Series B, Biological Sciences, 372(1713). 10.1098/rstb.2015.0484

MacConaill, L. E., Burns, R. T., Nag, A., Coleman, H. A., Slevin, M. K., Giorda, K., Light, M., Lai, K., Jarosz, M., McNeill, M. S., Ducar, M. D., Meyerson, M., & Thorner, A. R. (2018). Unique, dual-indexed sequencing adapters with UMIs effectively eliminate index cross-talk and significantly improve sensitivity of massively parallel sequencing. BMC Genomics, 19(1), 30.

Marciniak, S., & Perry, G. H. (2017). Harnessing ancient genomes to study the history of human adaptation. Nature Reviews. Genetics, 18(11), 659–674.

Mathieson, I., Lazaridis, I., Rohland, N., Mallick, S., Patterson, N., Roodenberg, S. A., Harney, E., Stewardson, K., Fernandes, D., & Novak, M. (2015). Genome-wide patterns of selection in 230 ancient Eurasians. Nature, 528(7583), 499–503.

Meyerhans, A., Vartanian, J. P., & Wain-Hobson, S. (1990). DNA recombination during PCR. Nucleic Acids Research, 18(7), 1687–1691.

Meyer, M., & Kircher, M. (2010). Illumina sequencing library preparation for highly multiplexed target capture and sequencing. Cold Spring Harbor Protocols, 2010(6), db.prot5448.

Mitra, A., Skrzypczak, M., Ginalski, K., & Rowicka, M. (2015). Strategies for achieving high sequencing accuracy for low diversity samples and avoiding sample bleeding using illumina platform. PloS One, 10(4), e0120520.

Neukamm, J., Peltzer, A., & Nieselt, K. (2021). DamageProfiler: fast damage pattern calculation for ancient DNA. Bioinformatics , 37(20), 3652–3653.

Olalde, I., & Posth, C. (2020). Latest trends in archaeogenetic research of west Eurasians. Current Opinion in Genetics & Development, 62, 36–43.

Oliva, A., Tobler, R., Cooper, A., Llamas, B., & Souilmi, Y. (2021). Systematic benchmark of ancient DNA read mapping. Briefings in Bioinformatics, 22(5). 10.1093/bib/bbab076

Patterson, N., Moorjani, P., Luo, Y., Mallick, S., Rohland, N., Zhan, Y., Genschoreck, T., Webster, T., & Reich, D. (2012). Ancient admixture in human history. Genetics, 192(3), 1065–1093.

Racimo, F., Renaud, G., & Slatkin, M. (2016). Joint Estimation of Contamination, Error and Demography for Nuclear DNA from Ancient Humans. PLoS Genetics, 12(4), e1005972.

Rasmussen, M., Guo, X., Wang, Y., Lohmueller, K. E., Rasmussen, S., Albrechtsen, A., Skotte, L., Lindgreen, S., Metspalu, M., Jombart, T., Kivisild, T., Zhai, W., Eriksson, A., Manica, A., Orlando, L., De La Vega, F. M., Tridico, S., Metspalu, E., Nielsen, K., … Willerslev, E. (2011). An Aboriginal Australian genome reveals separate human dispersals into Asia. Science, 334(6052), 94–98.

Rohland, N., Harney, E., Mallick, S., Nordenfelt, S., & Reich, D. (2015). Partial uracil-DNA-glycosylase treatment for screening of ancient DNA. *Philosophical Transactions of the Royal Society of London. Series B*, Biological Sciences, 370(1660), 20130624.

Rohland, N., Mallick, S., Mah, M., Maier, R., Patterson, N., & Reich, D. (2022). Three assays for in-solution enrichment of ancient human DNA at more than a million SNPs. Genome Research. 10.1101/gr.276728.122

Sinha, R., Stanley, G., Gulati, G. S., Ezran, C., Travaglini, K. J., Wei, E., Chan, C. K. F., Nabhan, A. N., Su, T., Morganti, R. M., Conley, S. D., Chaib, H., Red-Horse, K., Longaker, M. T., Snyder, M. P., Krasnow, M. A., & Weissman, I. L. (2017). Index switching causes “spreading-of-signal” among multiplexed samples in Illumina HiSeq 4000 DNA sequencing. In bioRxiv (p. 125724). 10.1101/125724

Skoglund, P., & Mathieson, I. (2018). Ancient Genomics of Modern Humans: The First Decade. Annual Review of Genomics and Human Genetics, 19(1), 381–404.

Weissensteiner, H., Forer, L., Fendt, L., Kheirkhah, A., Salas, A., Kronenberg, F., & Schoenherr, S. (2021). Contamination detection in sequencing studies using the mitochondrial phylogeny. Genome Research, 31(2), 309–316.

