## Supplementary Figures for "Optimised in-solution enrichment of over a million ancient human SNPs"

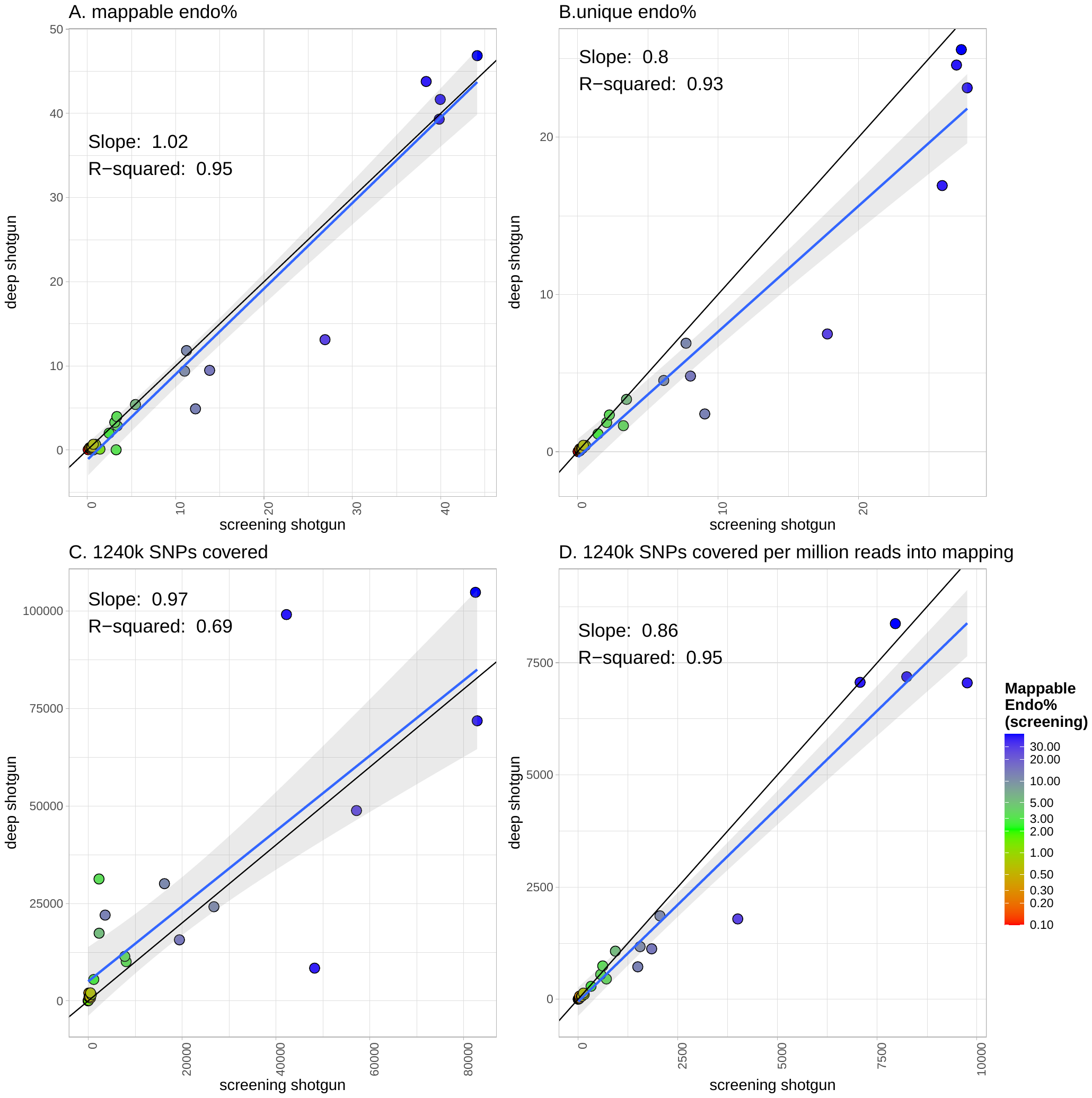

**Figure S1:** Comparative measures of library quality for shotgun screening data (x-axis) against deep shotgun sequencing (y-axis). (A) Mappable endo%, (B) unique endo%, (C) Target SNPs covered by at least one read, and (D) Target SNPs covered per million reads input to mapping. Black line shows y = x. For each plot, a linear model is fitted (blue) with a 95% confidence interval (grey shading), and the slope and *r*^2^ value is shown.

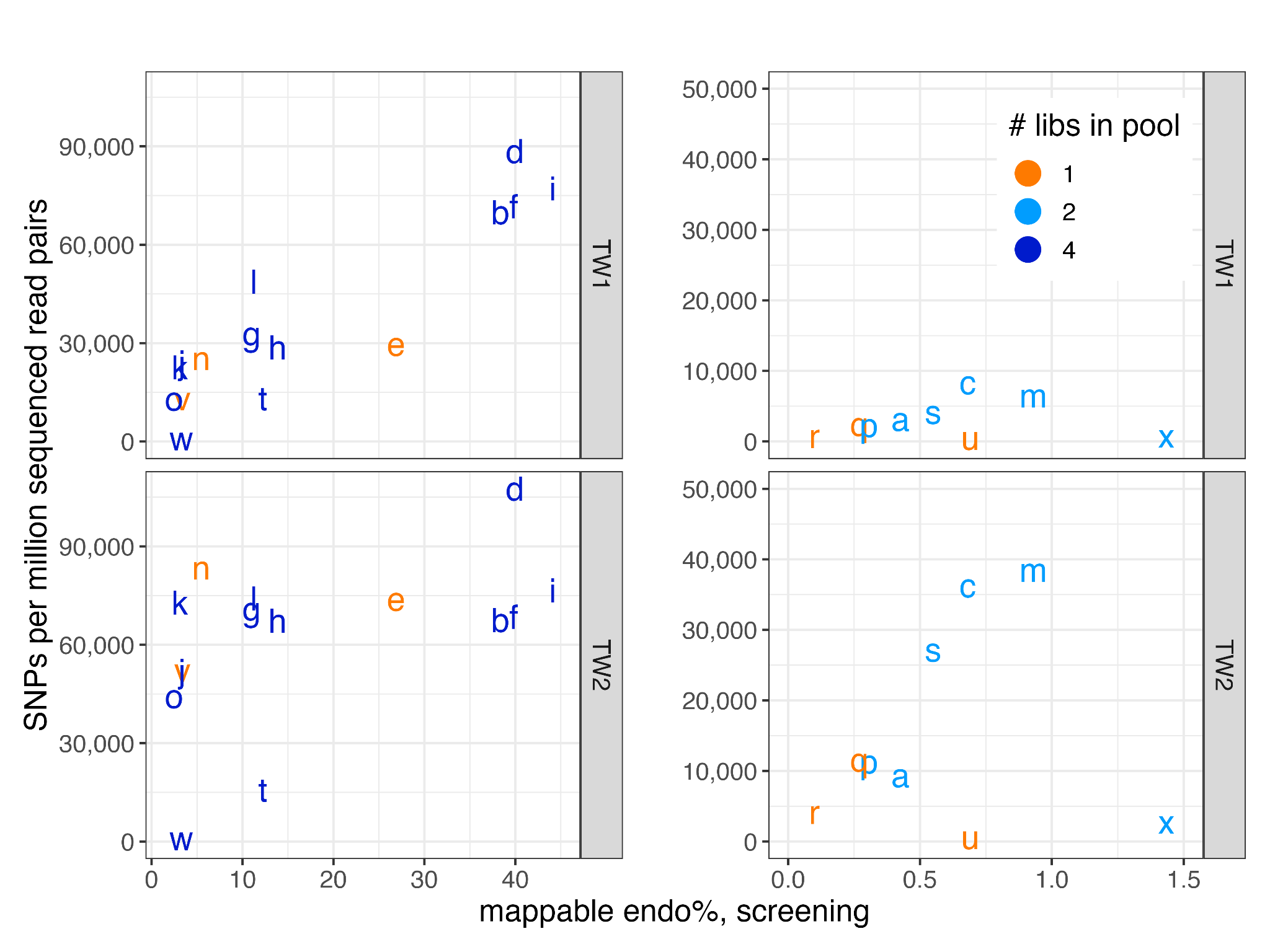

**Figure S2:** Number of target SNPs per million sequenced read pairs as a function of mappable endo% calculated from screening shotgun data. The different numbers of libraries in the pool are represented in different colours. Separated into TW1 (top) and TW2 (bottom), where the x-axis on the left is focussed on the range of 4-library pools: 1.5 - 45 endo%, and the x-axis on the right is focussed on the range of 2-library pools: 0 - 1.5 endo%

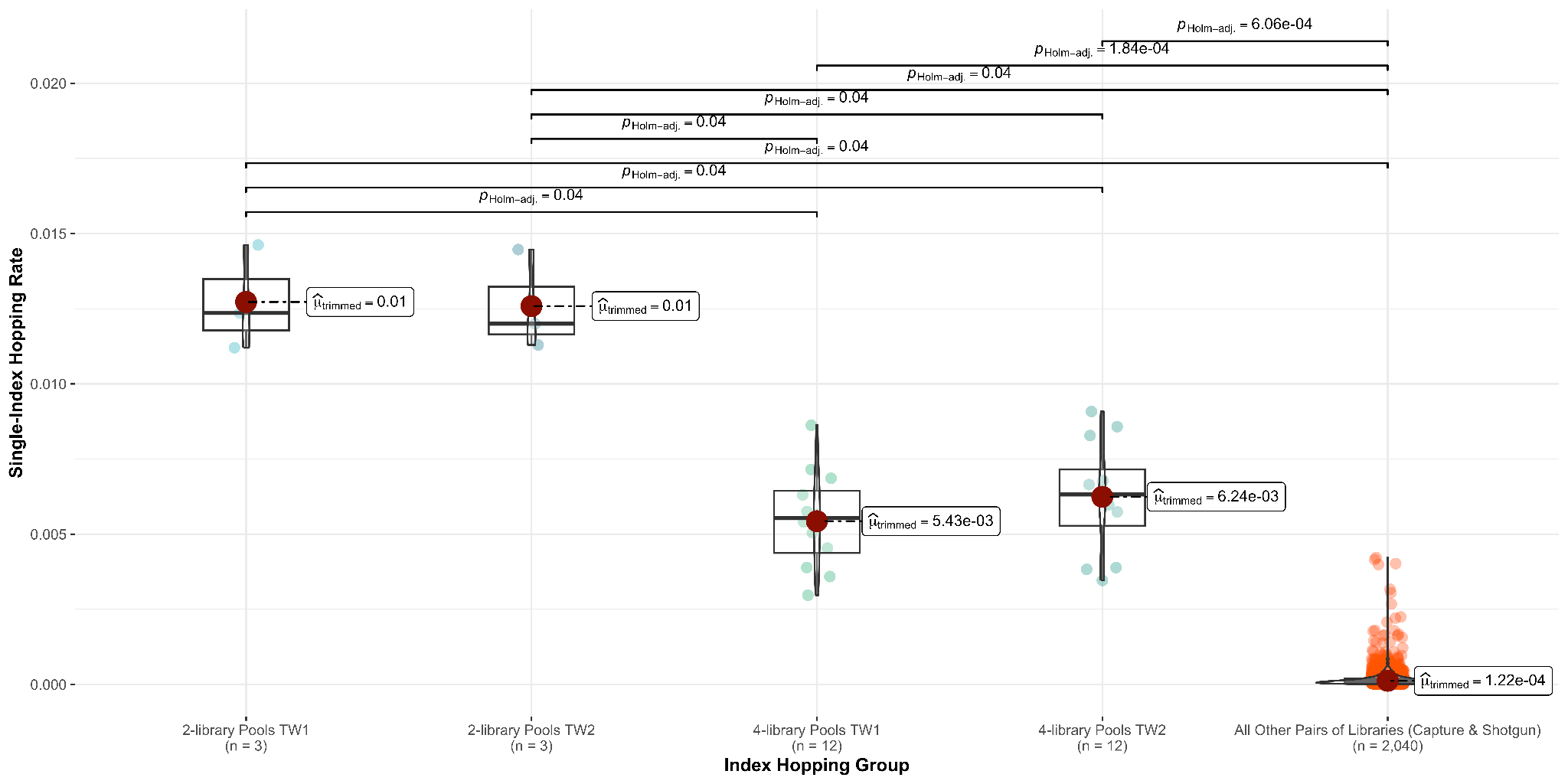

**Figure S3:** Pairwise rate of single index hopping between every pair of libraries, categorised into (left to right) 2-library pools TW1, 2-library pools TW2, 4-library pools TW1, 4-library pools TW2 and all other pairs of libraries. Each category is annotated with the number of pairs (*n*). The pairwise hopping rate was calculated as the proportion of reads with a hopped index combination over the sum of reads from every possible index combination in the pair.

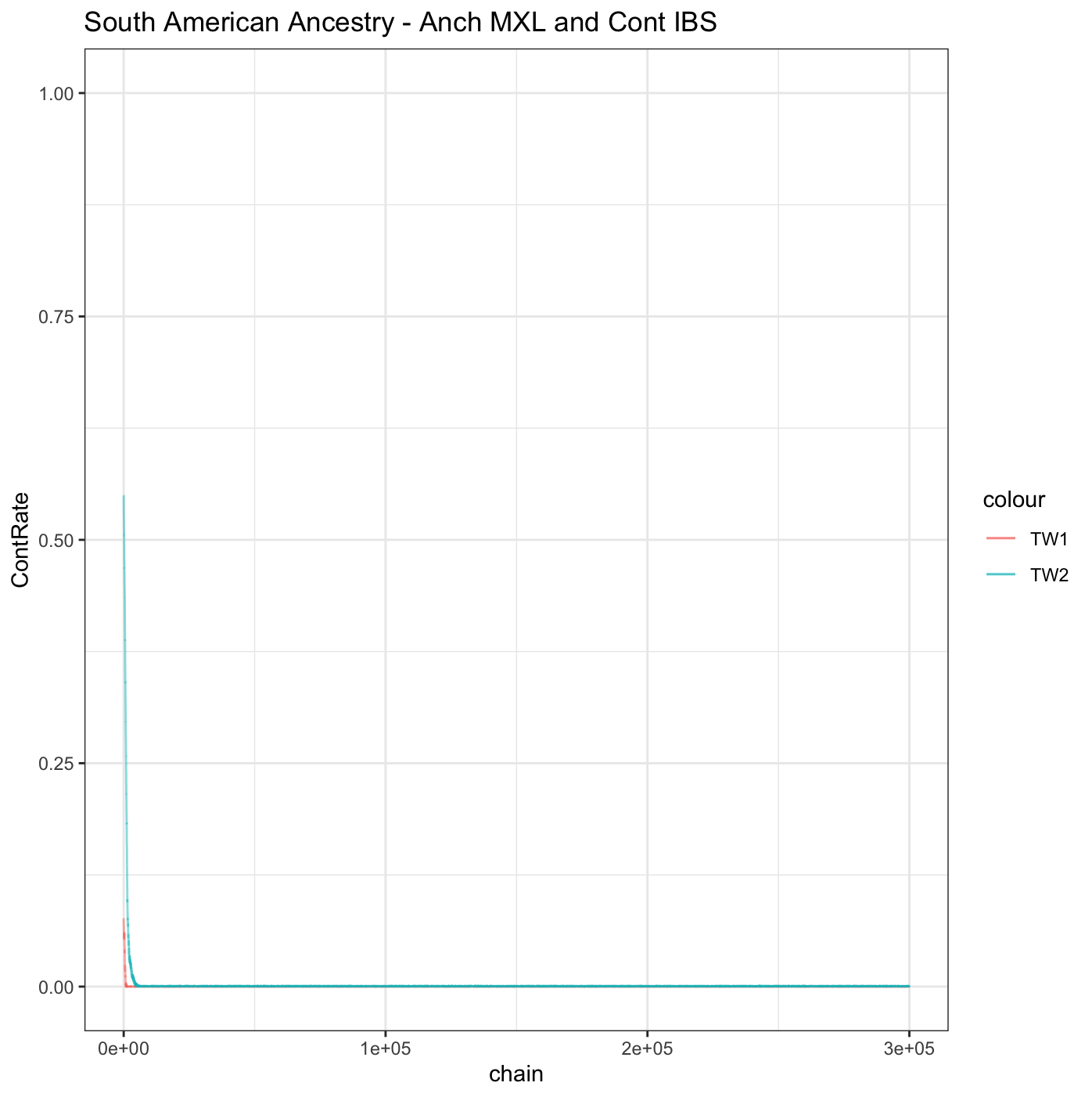

**Figure S4:** DICE contamination estimate for 300,000 MCMCs shows the contamination estimate converging towards 4.547888e-05 (SD = 3.546582e-05) and 0.0004988741 (SD = 0.0001391556) for Twist 1 and 2 round captures respectively.
